# Unpicking the mysterious symbiosis of *Mycoplasma* in salmonids

**DOI:** 10.1101/2020.07.17.209767

**Authors:** B Cheaib, P Yang, R Kazlauskaite, E Lindsay, C Heys, M De Noa, Patrick Schaal, T Dwyer, W Sloan, UZ Ijaz, MS Llewellyn

## Abstract

Lacking a peptidoglycan cell wall, mycoplasmas are the smallest self-replicating life forms. Members of this bacterial genus are known to parasitise a wide array of metazoans including vertebrates. Whilst much research has been significant targeted at parasitic mammalian mycoplasmas, very little is known about their role in other vertebrates. In the current study, we aim to explore the biology and evolution of *Mycoplasma* in salmonids, including cellular niche, genome size structure and gene content. Using Fluorescence *in-situ* hybridisation (FISH), *mycoplasmas* were identified in epithelial tissues across the digestive tract (stomach, pyloric caecum and midgut) during the developmental stages (eggs, parr, subadult) of farmed Atlantic salmon (*Salmo salar*), showing a high abundance in acidic compartments. With high throughput sequencing from subadults farmed Atlantic salmon, we assembled a nearly complete genome (~0.57 MB) via shotgun-metagenomics. The phylogenetic inference from the recovered genome revealed successful taxonomic proximity to *Mycoplasma penetrans* (~1.36 Mb) from the recovered genome. Although, no significant correlation between genome size and its phylogeny was observed, we recovered functional signatures, especially, riboflavin encoding genes pathway and sugars transporters, suggesting a symbiotic relationship between *Mycoplasma* and the host. Though 247 strains of *Mycoplasma* are available in public databases, to the best of our knowledge, this is the first study to demonstrate ecological and functional association between *Mycoplasma* and *Salmo salar* which delineates symbiotic reductive evolution and genome erosion primarily and also serves as a proxy for salmonid health in aquaculture processes (cell lines, *in vitro* gut models).

## Introduction

Commensal associations between bacteria and metazoan hosts are already well-established as being ubiquitous in nature. The extent to which microbes have adapted to persist in the intra-host environment varies considerably between taxa: some are opportunistic commensals, with others being obligate parasites or symbionts (1). Mycoplasmas are a diverse group of bacteria that are known to parasitise a wide array of metazoans, plants, invertebrates and vertebrates, including fish (1). In vertebrates, the mucosal surfaces of the alimentary canal, respiratory and genital tract are the primary site of colonisation (2). Mycoplasma *sp*. are a source of human and mammalian diseases. Of particular interest, but not limited to, is their implication in immunocompromised human cohorts (3, 4). It is generally thought that mycoplasmas have strict host associations, resulting in low zoonotic potential (5–7). Whilst there has been significant research effort targeted at parasitic mammalian *Mycoplasma* species, less is known about their importance and role in other vertebrates. Mycoplasmas, as well as related taxa included in the class Mollicutes (*Spiroplasmas*, *Ureaplasma* and *Acholeplasmas*), are recognized as the smallest and simplest free-living and self-replicating forms of life (6, 8). Mycoplasmas lack a peptidoglycan cell wall and are bounded by a simple cell membrane (7). In addition to being physically small, mycoplasmas have the smallest genomes of any free-living organism(2). *Mycoplasma genitalium*, in particular, has a genome size of 580 kilobases comprising of only 482 protein-coding genes(9), whilst *Mycoplasma mycoides*, typically has 473 protein-coding genes, of which 149 still have no known function(9). The relative simplicity of *Mycoplasma* genomic contents and structure has made this genus the target of scientific community’s efforts to design and synthesize a minimal bacterial genome, *de novo*, to establish the minimum requirements for biological life(10).

To further support this argument, the small size and simplicity of mycoplasmas, as well as their close association with metazoan hosts and their ability to survive as free-living species, irrespective of host, has led them to be considered as a target species to explore genome erosion or reductive evolution (11, 12) which refers to genes loss from an organism’s genome. Dependence on host organisms can theoretically lead to mutual interdependence of metabolic processes. This results in its relaxed selection amongst the pool of bacterial genomes, with the main process being the accumulation of loss-of-function mutations in coding genes, and the eventual loss of genetic material from the bacterial genome (13). Genetic drift can also play a significant role as host-associated microbes have relatively fewer opportunities to exchange genetic material (14). Enhanced mutational pressure from impaired DNA repair machinery could also be a factor (15). Isolation from microbial congeners and host dependence may be further enhanced in mycoplasmas that exploit an intracellular niche, which several species have been show to do within the literature (2, 16). *Mycoplasma penetrans*, for example, is predominantly important because of its ability to penetrate the host cells via an organelle specialised for host cell adherence (17). Mycoplasmas, likely owing to their dependence on their hosts, have fastidious requirements for *in vitro* culture. Culture-free approaches for microbial identification, especially, with the advent of DNA sequencing approaches, have markedly increased in the recent years to identify new *Mycoplasma*-like organisms(18–21).

Several studies have identified *Mycoplasma* from marine teleosts using culture-free approaches. Mudsucker (*Gillichthys mirabilis*) and pinfish (*Lagodon rhomboids),* for example, have been identified as having gut microbiomes rich in *Mycoplasma* (22). However, salmonids in particular are frequently reported to be colonised by *Mycoplasma* (23, 24). This is especially true in Atlantic salmon (*Salmo salar*), both in wild and in farmed settings and in farmed settings (23, 25). In some cases, *Mycoplasma* phylotypes can comprise >70% of the total microbial reads recovered from salmon intestines (24, 26).The distribution and biological role of *Mycoplasma* in the intestines of salmonids is far from clear, and requires further exploration. Nonetheless, demographic modelling of microbial communities suggest colonisation of salmonid guts by these organism as *non-neutral*, i.e. the rate at which these organisms colonise the gut, indicates a significant degree of specific adaptation to the host environment (26, 27).

In the current study, we aimed to explore the characteristics of *Mycoplasma* in salmonids, including cellular niche, taxonomic affiliations, genome structure and gene content. We focused on the genetic features and metabolic functions which may help us to explain the role of reductive evolution in the close association of the *Mycoplasma* with the host, through different physiological and physio-chemical adaptations to survival within the digestive tract of *Salmo salar*. We also explored the phylogenetic relatedness of the *Mycoplasma* in salmonids as compared to all the known and sequenced mycoplasmas to date.

## Materials and Methods

### Sample collection

Farmed *Atlantic salmon* (*Salmo salar*) subadults (3 to 5 kg) were sampled from marine cages at an aquaculture facility at Corran Ferry, near Fort William, Scotland, in Autumn 2017 in collaborations with MOWI Ltd. *Salmo salar* freshwater parr and ova were sampled at the Institute of Biodiversity, Animal Health and Comparative Medicine aquarium facility, University of Glasgow. Animals were euthanised by blunt cranial trauma under a Schedule 1 procedure and gut compartments (stomach, pyloric caecum, and midgut) samples were flash frozen in liquid nitrogen and stored in −80 C.

### Fluorescence *in-situ* hybridisation (FISH)

Previous work has established the dominance of *Mycoplasma* in marine *Salmo salar* GI (gastrointestinal tract) (26). To explore their physical distribution in different gut compartments and life cycle stages, FISH was undertaken on salmon tissues. Samples were fixed in a freshly made sterile-filtered solution of 4% paraformaldehyde in PBS (pH 7.4) for 16-24 hours and maintained at room temperature for 16-48 hours. Fixed samples were then washed with sterile-filtered PBS (pH 7.4) three times before being fixed the sample in 70% ethanol. Samples were then gradually dehydrated in a series of ethanol-xylene-paraffin treatment steps (28). Prior to sectioning, samples were embedded in paraffin and stored at 4℃. At least four 3-4 µm sections were taken from each embedded tissue block, rehydrated in sterile ddH_2_0, and mounted on slides for pepsin treatment and straining. Pepsin treatment was undertaken in a 0.05% pepsin solution and 0.01M HCL. Samples were DAPI stained to target cell nuclei of host cells, and FISH probes were hybridised at 55°C to the 16S rDNA small subunit of bacterial cells. Multiple FISH probes labelled with Cy3 and Cy5 dyes were deployed to distinguish *Mycoplasma* from other microbes present in samples (**Table 1**). To improve the visualisation of non-*Mycoplasma* bacteria, multiple probes were deployed using the same dye. A *Mycoplasma* (Myc1-1, **Table 1**) probe was designed based on Illumina amplicon sequences based upon the most abundant operational taxonomic (OTU) sequence identified in Adult Salmon that we identified in previous work (Heys, Cheaib et al 2020). Samples were visualised at 20-30x magnification on a DeltaVision-Core microscope (Applied Precision, GE), equipped with a CoolSNAP HQ camera (Photometrics) and operated with SoftWoRx software (Applied Precision, GE).

**Table 1.**
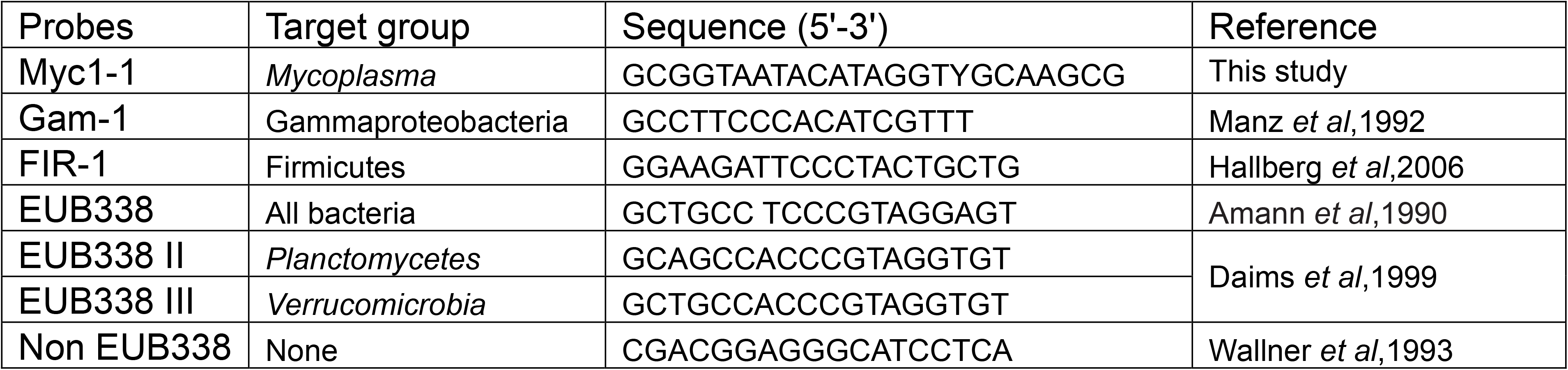
Summary of FISH probes used in this study

### DNA extraction, library annotation and sequencing

DNA was extracted from the pyloric caecum homogenate derived from a single individual on which FISH analyses had identified the presence of *Mycoplasma*-like organisms, based on their labelling with a targeted 16S probe. A sequence library for Illumina Next-Seq WGS (whole genome shotgun) sequencing was prepared using a sonication protocol and a TruSeq library protocol and sequencing adaptors. Sequencing was undertaken at the University of Glasgow Polyomics facility.

### Data preprocessing, assembly, binning and annotations

The short paired-end NextSeq Illumina reads (2 × 63 million reads) were preprocessed for quality filtering using sickle V1.2 (https://github.com/najoshi/sickle). Decontamination of good quality reads was performed by mapping reads against the *Salmo salar* genome (available at NCBI sequence archive with the accession number GCF_000233375.1) using Deconseq V 0.4.3 (29) based on BWA mapper V 0.5.9 (30). The decontaminated paired-end reads (~18 million of bacterial reads) were assembled using the Megahit V1.1 software (31). The assembled contigs (~93400) were processed for genomic binning using MetaBAT V2.12.1 (32). Quality assessment for completeness and contamination of sequence bins was performed using CheckM V1.0.18 software (33). Annotation of gene content was performed using the pipeline ATLAS-metagenome (34), which involves the prediction of open reading frames (ORFs) using Prodigal (35). Translated gene products were clustered using linclust (36) to generate non-redundant gene and protein catalogues, which were mapped to the eggNOG catalogue (37) using DIAMOND (38).

### Phylogenetic analyses

Two approaches were undertaken to construct phylogenetic trees: a) MLST-based (Multi Locus Sequence Typing); and b) 16S gene markers (recovered from the genome) of the *Mycoplasma* MAG (metagenome assembled genome) from this study as well as what is previously available in the literature. Using CheckM software, the MLST-based strategy focused on a concatenation of 21 conserved housekeeping genes annotated in the *Mycoplasma* bin supplemented with the orthologues available for all the *Mycoplasma* genera, to date. The MLST-based dataset included 55 orthologues of protein sequences of concatenated 21 markers. The 16S rDNA sequence dataset included: one sequence of 16s annotated in the *Mycoplasma* bin; five Operational Taxonomic Units (OTUs) sequences of mycoplasmas characterised form the same farmed salmon system (26); 101 and 17 sequences of 16S rDNA from the *Mycoplasma* and *Sprioplasma* genomes respectively from IMG database; and 11 sequences from environmental studies detected in marine species including shrimp, fish and isopods. DNA and protein sequences were aligned using MAFFT version 6.24 (39). Phylogenetic inference was performed using PhyML version 3.0 (40) and MrBayes V.3.2.6 (41). The evolutionary model was chosen using MODELTEST(42), and parameters were iteratively estimated in PhyML using the GTR+I+G model for nucleotide sequence of 16s trees and the LG+I+G model for amino-acid sequence of concatenated markers trees (43). Bootstrap values were calculated with 100 replicates (44). With MrBayes, posterior probability values were calculated using an average standard deviation of partition frequencies < 0.01 as a convergence diagnostic (45). MrBayes runs consisted of eight simultaneous Markov chains, each with 1,000,000 generations, a subsampling frequency of 1000, and a burn-in fraction of 0.15. Trees were then visualized and adapted for presentation in FigTree version1.4.3 as a graphical viewer of phylogenetic trees (http://tree.bio.ed.ac.uk).

### Metabolic pathways comparison and genome reduction analysis

All pFam V.32 (comprehensive and accurate collection of protein domains and families) annotations were predicted with Prodigal and analysed in terms of function categories and metabolic content (focusing on Enzyme EC numbers). The 530 genes identified were associated with 746 pFam functions. The pFam function led to the recovery of Gene Ontology (GO) terms and were then mapped to the KEGG database. Simultaneously, the alternate approach involving MetaCyc database was employed to elucidate metabolic pathways from all domains of life (46). The EC numbers of the coding sequence regions in *Mycoplasma penetrans* was extracted from the KEGG database and was then compared with those annotated within the MAG of *Mycoplasma* from *Salmo salar* in this study. The mapping of metabolic pathways from both genomes were visualized using the iPath (47). From the IMG genomic database, all available metadata on sequenced *Mycoplasma* strains were then collected and compared to the MAG for the genome size, GC content, gene content and their preference (e.g. intracellular, free-living etc). Annotations for the assembled *Mycoplasma* genome were submitted to CG view (48) for radial visualisation of genomic structure of the assembled *Mycoplasma* bin. The 570 predicted genes were compared at the DNA and protein sequence levels against all the available genes of *Mycoplasma penetrans* using BLAST+ V 2.8.1 (49). Best hits for each query were represented in a radial plot using Circoletto software version V.069-9 (50). Complimentary annotations were performed using RAST software which, consisted of subsystem classification of microbial functions available in the curated database, i.e. SEED subsystem (51).

## Results

### Fluorescence in situ hybridization (FISH) of mycoplasmas in the farmed salmon

The set of probes used in FISH for the identification of bacterial populations is summarized in the **Table 1**. Only the Myc1-1 probe was shown to be specific to *mycoplasmas* (**Supp. Figure S1**). FISH visualization in salmon ova demonstrated low abundance of bacteria and no signal of mycoplasmas (Figure 1-a; **Supp. Figure 2.1**). In *Salmo salar* freshwater parr, *Mycoplasma* aggregates were observed on the stomach lining (Figure 1-b; **Supp. Figure 2.2**), as well as on the muscularis mucosae, and epithelium of the pyloric caecum (**Figure 1-c**; **Supp. Figure 2.3**). In the midgut of salmon parr (distal to the pyloric caecum) we found some evidence that the *Mycoplasma* may be clustering intracellularly (**Supp. Figure 2.4**). In the stomach (Figure 1-d; **Supp. Figure 2.5**) and pyloric caecum (**Figure 1-e-f**; **Supp. Figure 2.6**) of adult salmon, *Mycoplasma* signals were clearly clustered in small aggregates in the lumen around the nuclei of epithelial cells. Figure 1 indicates this intracellular clustering most clearly. In the midgut of adult salmon, *Mycoplasma* showed low abundance and the signals of *Mycoplasma* showed aggregations near epithelium cell nuclei (**Supp. Fig 2.7**). The specificity of the *Mycoplasma* probe against pure culture *Escherichia coli* and *Mycoplasma Muris* was evaluated (**Supp. Figure S1**). The *Mycoplasma* probe Myc1-1 showed specific hybridization, giving a positive signal solely with cultured *Mycoplasma muris*. We made multiple attempts in both solid and liquid culture mycoplasmas from the salmon intestines, but without success.

**Figure 1.**
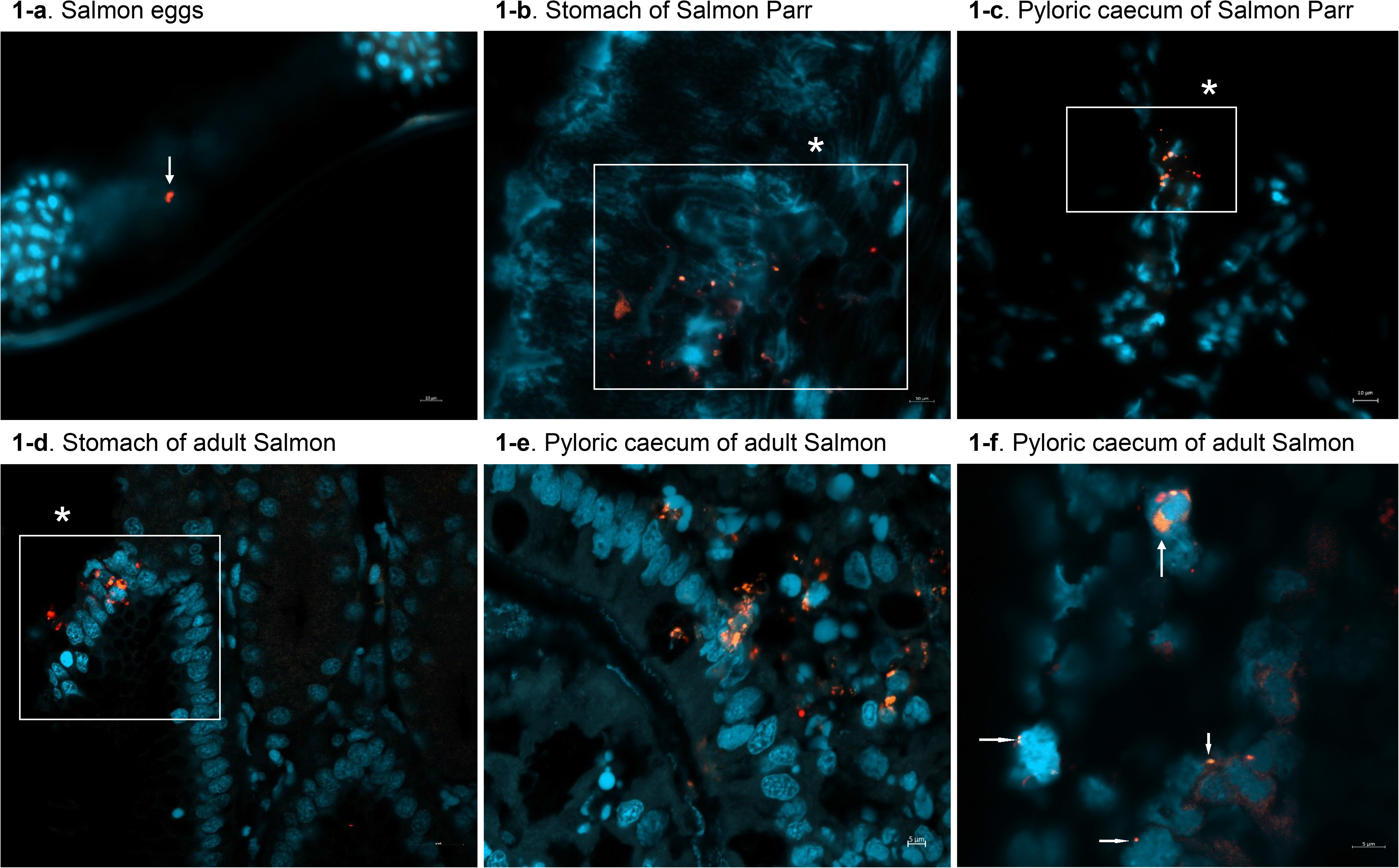
FISH visualization of *Mycoplasma* in acidic gastrointestinal tracts of salmon Parr and adults. The images were an overlay of DAPI signals (blue), hybridization signals of Gam-1, FIR-1, EUB338, EUB338 II, EUB338 III probes (Cy5, red) and *Mycoplasma* Specific Myc1-1 probe (Cy3, orange). (A) Low *Mycoplasma* distribution in Salmon egg, the image is an overlay of Cy3, Cy5, and DAPI filter set showed bright signal scaled at 10 μm.(B) Distribution of *Mycoplasma* in the stomach of Salmon Parr, scaled at 10 μm. Orange signals indicate mycoplasmas were clustered in small groups. (C) Distribution of *Mycoplasma* in the epithelium of Pyloric caecum of Salmon Parr at 10 μm. (D) Distribution of *Mycoplasma* in Stomach of adult Salmon scaled at 50 μm. (E, F) Distribution of *Mycoplasma* in Pyloric caecum of adult salmon scaled at 10 μm (E) and 5μm (F) respectively. Mycoplasmas signals were aggregated on the muscularis mucosae, lamina propria (E) and clustered in high abundance around epithelial cells nuclei (F). (*) indicatesa is frame of micrograph detail. Horizontal bars indicate that these images are scaled at 10 μm for (A, B, C, D, E), and 5 μm for (F).

### Binned *Mycoplasma* genome features and orthologs

Using a total of 63,180,207 reads, and after decontamination, 93397 contigs were assembled using megahit software (see materials and methods). The assembled contigs were binned, annotated and assessed for completeness (see materials and methods). The best quality assembled bins corresponded to a nearly complete genome assigned to *Mycoplasma* (see bin sequences in **Supp. File 1**). The completeness of this metagenome-assembled genome (MAG) was estimated at 98 % with 0.38 % of contamination and 0.0% of heterogeneity (**Table 2)**.

**Table 2.** Summary of *mycoplasma* metagenome assembled genome (MAG) features.

|  |  |
| --- | --- |
| Genome | *Mycoplasma* bin (TMDNneg-S20-001) detected in Adult Salmon<br>*Salmo Salar* |
| Completeness | 92.18 |
| Contamination | 0.38 |
| Strain heterogeneity | 0 |
| Unique markers (of 43) | 39 |
| Multi-copy | 0 |
| Taxonomy (contained) | k__Bacteria; p__Tenericutes ;c__Mollicutes;<br>o__Mycoplasmatales; f__Mycoplasmataceae ;g__Mycoplasma |
| Taxonomy of sister lineage | s__Mycoplasma_penetrans |
| GC content | 24.98204716 |
| Genome size (mbp) | 0.577903 |
| Gene count | 570 |
| Coding density | 0.938088226 |
| Length | 577903 |
| N50 | 14796 |
| Comment 1 | Genome with >90% completeness and < 5% contamination |
| Comment 2 | Binned with Metabat and quality assessed with CheckM |

The average size of the assembled genome was estimated to be 0.557 Mb and comprised a set of 570 predicted genes accounting for a total of 694 CDS regions found on the 5’3’ and 3’5’ ORFs (supplementary file 2). The GC percentage was estimated to be 39.2% **(Table 2).** Circular representation of the genomic structure of the MAG highlights CDS annotations (694) on the negative (**Figure S3-a**) and positive (**Figure S3-b**) strands, respectively. To further resolve CDS annotations, a supplementary annotation framework was applied using the curated SEED database and the RAST server (52). The results showed 600 CDS across the negative (275 CDS) and positive strands (325 CDS). Amongst these CDS regions (**Supp. File 2**), 390 had functional annotations, and within these, three annotated CDS regions (> 85% of similarity threshold against SEED) were identified as: Riboflavin kinase (EC 2.7.1.26/EC 2.7.1.26; 1278 bp) along with two Riboflavin/purine transporters of length 1383 bp and 1608 bp, respectively. In addition, other functions required for host-microbiota symbiosis, such as ribonucleotide reductase, were annotated with SEED and are reported (**Supp. Table 1**).

### Phylogenetic proximity to *Mycoplasma penetrans*

The recovered phylogenetic tree based on the MLST approach as well as 16S rDNA, corroborated the same genetic relatedness of the assembled *Mycoplasma* to the closest lineage represented by *Mycoplasma penetrans.* The 16S rDNA tree includes four OTUs of *Mycoplasma* detected in the digestive tract of farmed salmon previously (26). The phylogenetic distances indicate that the *Mycoplasma* MAG is closer to the two OTUs from the same farmed system as compared to *M. penetrans* (**see sequence alignment in Supp. File 3)**. Furthermore, clusters containing these taxa are supported with medium to high posterior probabilities (>0.5) according to the Bayesian approach (**Figure 2).**

**Figure 2.**
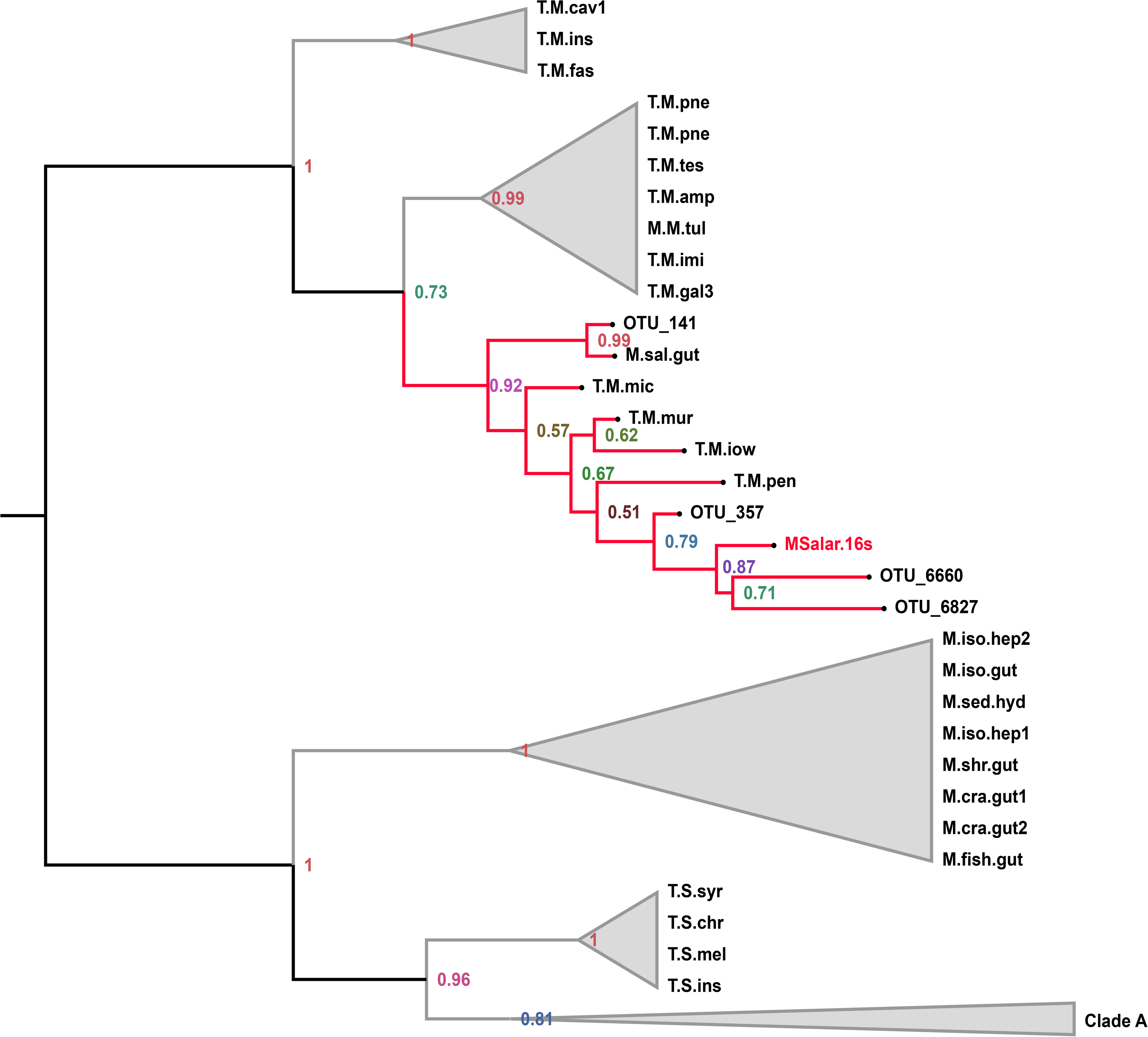
Phylogenetic tree based on 16S rRNA gene sequences of *mycoplasmas.* Sequence name abbreviation of tree tips labels and clade A (including spiroplasmas) are reported in Supplementary File 6.

To further ascertain the above clustering of 16S rDNA sequences of *Mycoplasma*, and the phylogenetic relatedness, a second analysis based on MLST approach using 21 concatenated housekeeping genes (see PFAM IDs of markers and their functions in Supp. **File 4**) increased our confidence in *M. penetrans* being close to the recovered *Mycoplasma* MAG (**Figure 3; see sequence alignment in Supp. File 5**). These 21 markers are detected in single copies and are conserved in the bacteria and the *Mycoplasma* lineage (). The MLST tree shows high posterior probabilities in support of this argument (post prob > 0.9). Tip labels of the selected *Mycoplasma* samples are further annotated with the genome size information in Mbp. It should be noted that the genome sizes did not appear to correlate well with the spatial distribution patterns of *Mycoplasma* species in the tree. The genome size of *M. penetrans* (1.36 Mb) is approximately double to that of the binned *Mycoplasma* and, is the highest amongst the *Mycoplasma* genomes.

**Figure 3.**
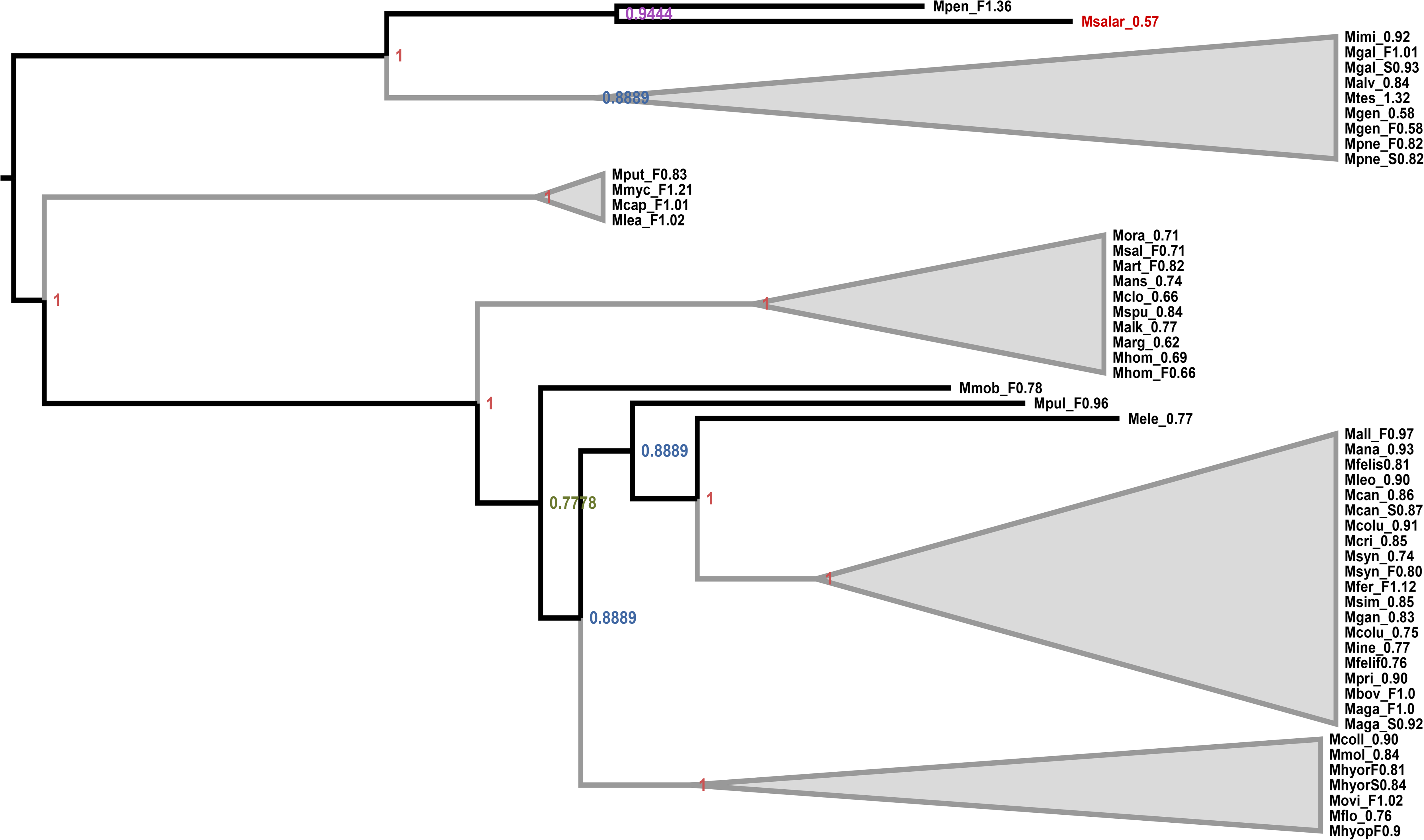
Phylogenetic tree of mycoplasmas based on 21 MLST markers with the details given in the supplementary data. Sequence name abbreviation of tree tips labels are explained in Supplementary File 7.

### Orthology, metabolic pathways and genome reduction analyses

A core genome analysis including the amino acid sequences of predicted CDS and all the available CDS from the closely related *M. penetrans*, available on NCBI repository, were blasted against the COG database. Circular track of the genome including the orthology clearly showed the difference in genome size between the binned *Mycoplasma* and *M. penetrans*. We observed heterogeneity across the different regions of the genomic structure in terms of GC content and GC skew (**Figure 4**). An orthology analysis based on SEED annotations indicated 14 functions (oxidative stress, periplasmic stress, protein biosynthesis, detoxification, ribonuclease H, cation transporters, ABC transporters) specific to the binned genome, 144 functions specific to *M. penetrans*, and 156 functions that are common to *Mycoplasma* MAG and *M. penetrans*. The shared functions between these two genomes belong to nine different general subsystems including those related to symbiosis and intracellular lifestyles such as: riboflavin metabolism; intracellular resistance; and resistance to antibiotics and toxic compounds (RATC) (**Supp. Table 2**). We only found two similarity hits associated to RATC. Complimentary analysis pointed out a bifunctional riboflavin kinase/FMN FMN adenylyltransferase among the best reciprocal similarity’s hits between the *Mycoplasma* MAG in this study and *Mycoplasma penetrans* (Figure 5; Supp. Table 3).

**Figure 4.**
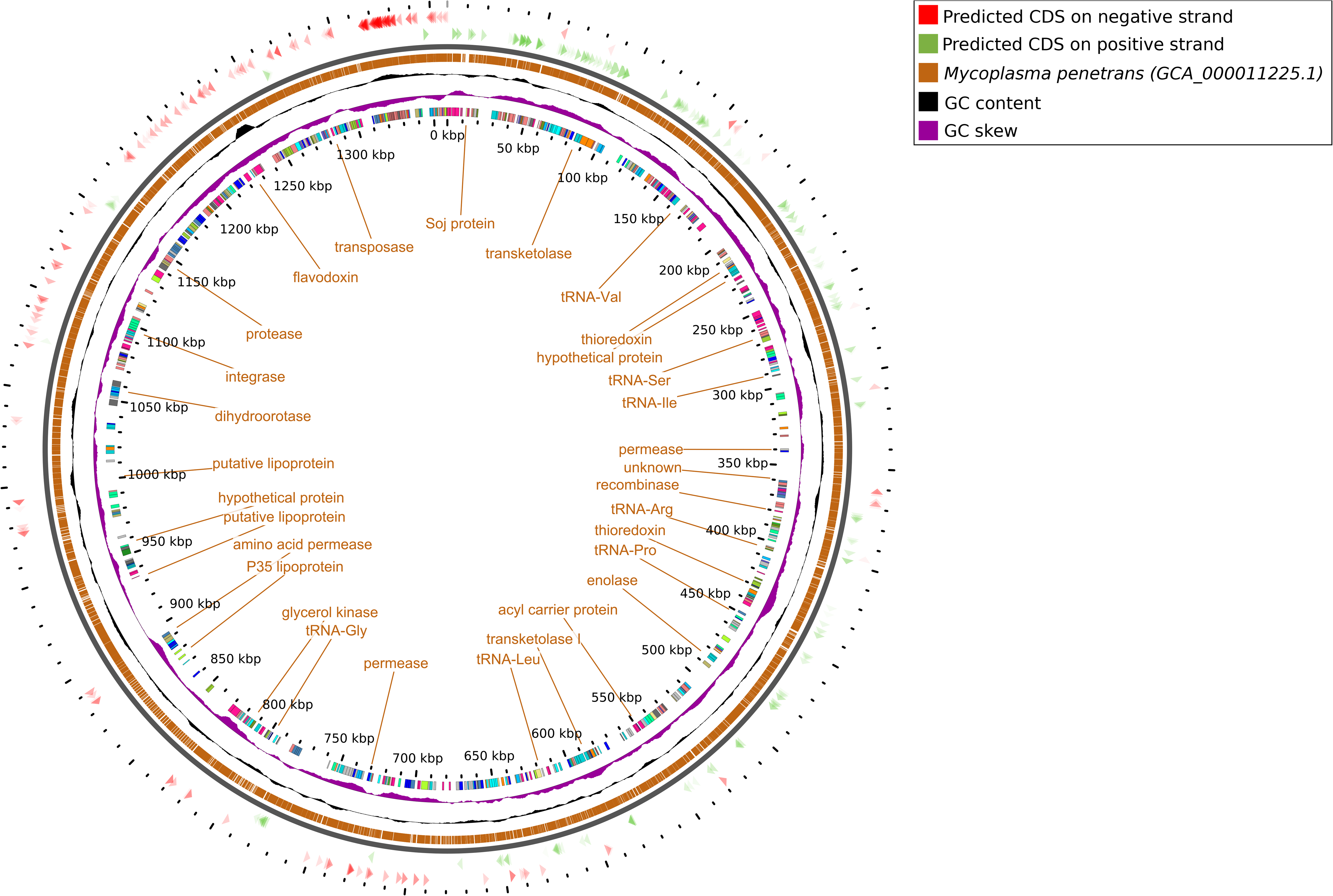
Circular track of the core genomes. This figure highlights core genes shared between the *Mycoplasma* MAG from this study and related *Mycoplasma penetrans* species.

**Figure 5.**
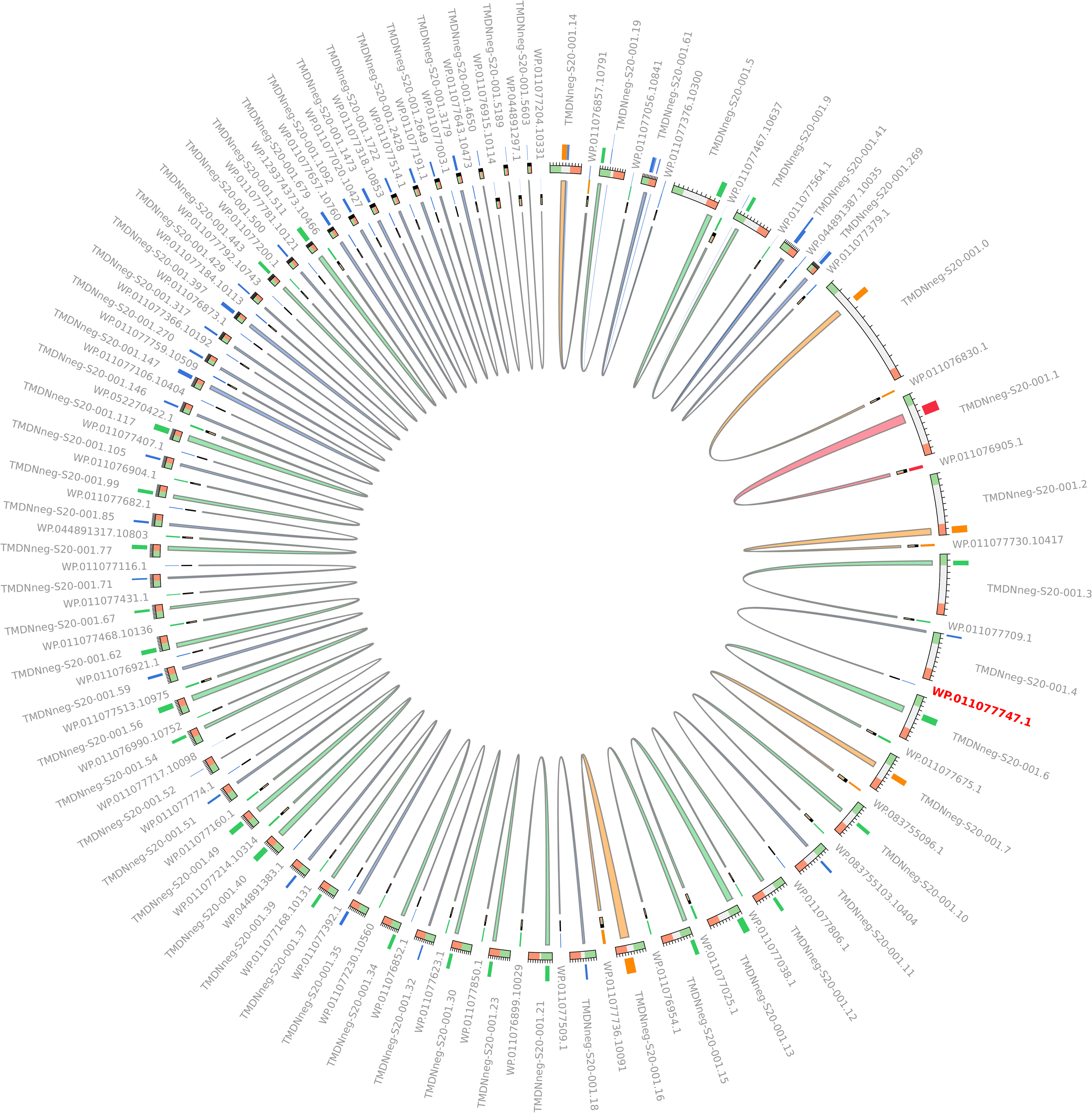
Circular track of best BLAST similarities. This figure highlights orthologous CDS regions between the *Mycoplasma* MAG from this study and *Mycoplasma Penetrans.* The orthologous CDS (WP.011077747.1) labelled in red encodes for bifunctional riboflavin kinase/FMN adenylyltransferase in *Mycoplasma penetrans*. See abbreviations of orthologs from this figure in Supp.Table 3.

To understand genome reduction in *Mycoplasma* lineage, the genome size and genes count were compared across 247 strains (**Figure 6-a; Supp. Table 4**) of the available *Mycoplasma* database (IMG) isolated from a wide variety of human and animal sources and comprising both parasitic and commensal strains. In view of the collected data (**Supp. Table 4**), gene content and genome size are strongly correlated. The average genome size of the 247 available mycoplasmas was 0.87 Mbp ± 0.15, and the average genes count was 790 ± 157 genes; however, this was not the case with all considered genomes. For instance, 8 genomes are lower than 0.8Mb, accumulating somewhere between 829 and 1036 genes. Further analysis revealed that pseudogenes count was not a significant factor, whilst both the transmembrane proteins and GC content were correlated with Mycoplasma genome sizes (**Figure 6-b).** Furthermore, the average count of pseudogenes was significantly higher in free-living than within intracellular mycoplasmas (**Supp. Figure S4**), although available databases contain incomplete information with regards to *Mycoplasma* lifestyles. Enzyme content was analyzed in terms of metabolic pathways by comparing the annotated EC numbers of the *Mycoplasma* MAG and *M. penetrans*. Common pathways of both genomes are highlighted in red lines (**Figure 7**) and include the riboflavin biosynthesis pathway.

**Figure 6.**
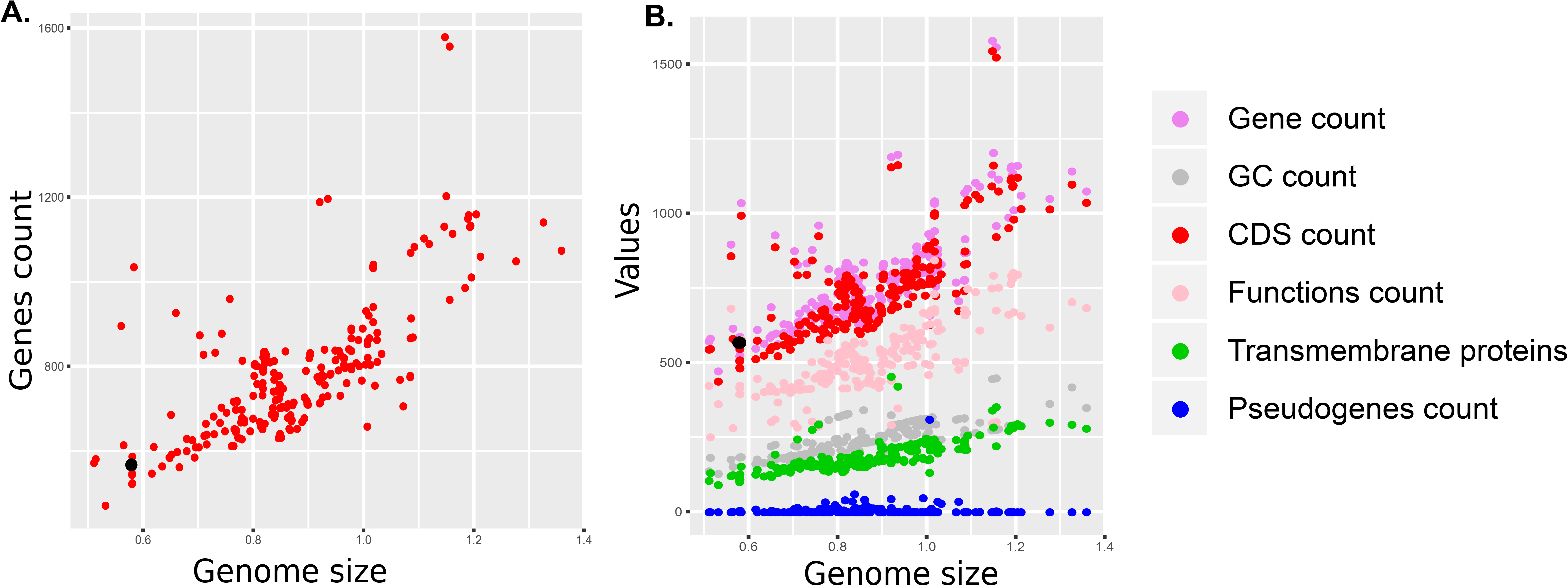
Genomic features of Mycoplasmas. (A) Plot of genome size and genes count in the Myoplsamas lineage. (B) Plot of functions and GC contents against genome size in the Mycoplasmas lineage

**Figure 7.**
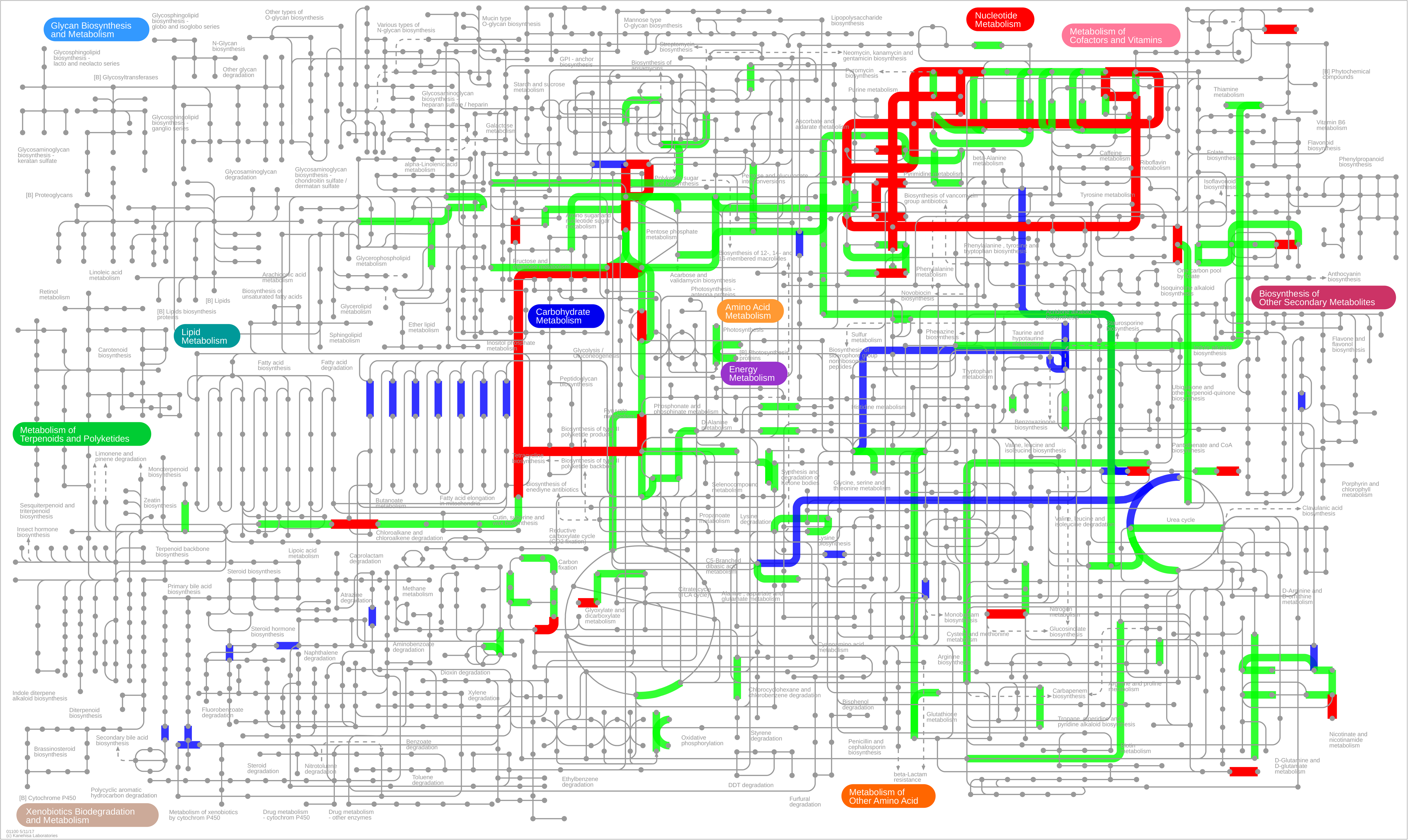
Pairwise Metabolic pathways comparison of *Mycoplasma* MAG from this study and related *Mycoplasma penetrans* species. Red color represents shared and conserved pathways between the two genomes, whereas blue color represents the metabolic pathways of the *Mycoplasma* MAG from this study and the green color represent the metabolic pathways of *Mycoplasma penetrans*.

## Discussion

*Mycoplasma* are hyper-abundant commensals of salmonid guts. Our study suggests, based on FISH data, that in *Salmo salar*, these organisms grow intracellularly in the epithelial and possibly muscular lining of the fish’s GI tract, both in freshwater and marine lifecycle stages. *Mycoplasma* sequences recovered from *Salmo salar*, including the *Mycoplasma* MAG reported here, had a strong phylogenetic closeness to *M. penetrans*. Comparative analysis of genome size and content across *Mycoplasma* strains suggest that the genome we recovered in this study is among the smallest ever observed, to the best of our knowledge. Comparative genomics analyses between the *Mycoplasma* MAG and *M. penetrans* were undertaken and provide insight into the potential host-microbe interaction.

Mycoplasmas have been widely reported within *Salmo salar* (23), and other teleosts (27). It is not uncommon to find that communities of gut microorganisms are dominated by mycoplasmas (53). The modelling approaches comparing environmental and intestinal frequency distributions of these organisms undertaken in this study have previously suggested that salmon mycoplasmas are well adapted to colonisation of their hosts (26). On the flip side, culture-based approaches have had been less successful in isolating these organisms (54) and despite numerous attempts, we failed to obtain pure cultures of *Mycoplasma* from the adult salmon used in this study (data not shown). This may be attributed to a potential source of bias arising from cell wall deficiency (55) in mycosplasmas which decrease their growth in presence of inhibitors such as nucleoside and nucleobase analogs as demonstrated in *Mycoplasma pneumoniae* (56) and others mycoplasmas (57). FISH data from the current study, however, indicate that many mycoplasmas are sequestered within the basal the epithelial cells, suggesting potential unknown parameters in symbiosis with *Salmo salar* which were missed from the culture media trials and reduce their cultivability. Although only a qualitative assessment is possible by employing FISH, our data suggest that *Mycoplasma* comprised the majority of the resident microbes (**Figure 2**), and the binding efficiency between *Mycoplasma* and the universal probes differed only slightly in the cultured controls (**Supp Figure S1**).

The intracellular niche of in the epithelial cells of the GI tract of salmon may represent a strategy to avoid hydrolysis in the intestinal environment. Sequestration in the muscularis under the basal of epithelium, as we observed in the stomach, is thought to offer a protective niche in the case of invading *Helicobacter pylori* in human stomach diseases (58). Also similar to *H. pylori,* our functional annotation of the *Mycoplasma* MAG (**Supp Table 1**) identified components of the urea cycle subsystem that may assist acid tolerance via the hydrolysis of urea to ammonia (59). Further investigations of gene complexes involved in acid neutralization in the *Mycoplasma* could potentially be achieved via targeted transcriptomics.

A high level of adaptation to, and dependence on the host organism, is a key feature of many *Mycoplasma* species (60). The exploitation of an intracellular niche, dependence on the host, and relative isolation from the other microorganisms and mobile genetic elements are thought to have contributed to genome decay in mycoplasmas (61). One result of this decay is a reduction in genome size and the number of genes, and the *Mycoplasma* MAG in this study appears to have been potentially affected by such processes in comparison to the other mycoplasmas (**Figure 5**). According the phylogenetic tree, we did not observe any specific relationship between the tree topology and the genome sizes of mycoplasmas (Figure 2). Indeed, the closely related *M. penetrans* was over three times larger than the size of the *Mycoplasma* MAG in this study. Despite sharing a recent ancestor with the human pathogen *penetrans*, a long and close evolutionary association of this *Mycoplasma* strain and salmonids is possible given the similarity of another *Mycoplasma* MAG sourced from the Norwegian sea salmon and identified to *M. penetrans* (62). We were also able to identify *Mycoplasma* in freshwater parr via the FISH method in this study. One potential route for vertical transmission of the *Mycoplasma* among salmon could be observed during oviposition. We were not able to identify microbes colonising eggs in this study, although our sample size was limited. Further development on specific *Mycoplasma* strain markers could potentially reveal their abundance as well as their epidemiology, and potential routes of inter and intra generational transmission.

Many well-characterised mycoplasmas are pathogens (17, 63, 64), with several Mycoplasma sp. being responsible for human, animal and plant diseases; however some species are considered to be commensal organisms (2, 65). The role of *Mycoplasma* in the context of *Salmo salar* is not well established. Koch’s postulates were not applied in this study (66). Given the challenges encountered in culturing these microorganisms, it seems quite likely that they may never be applied. Furthermore, the apparent abundance of *Mycoplasma* in the healthy salmonids (23, 24, 26), and lack of any clear associated pathology in gut tissues, implies that there is not a significant impact on the host health or fitness. Commensal exploitation of the host intracellular niche is potentially the most parsimonious description of the host-microbe interaction. The ultimate metabolic adaptation to an intracellular lifestyle (i.e *Buchnera*, *Wigglesworthia* and *Blochmannia)* appears to be solely regulated by the metabolic activity of the host cells to which the bacteria may actively contribute to, by delivering essential metabolites that are limited in their habitats and are not produced by the hosts (67). Nonetheless, the presence of an apparently complete riboflavin pathway could potentially indicate benefit from the host perspective of *Mycoplasma* colonisation. Riboflavin, known as the precursor for the cofactors flavin mononucleotide (FMN) and flavin adenine dinucleotide, is an essential metabolite in organisms (67, 68), although vertebrates cannot synthesise it on their own (69). The *Mycoplasma* likely plays a role in riboflavin supplementation in salmon, as observed in several deep-sea snailfish (70). Riboflavin supplementation is not limited to the *mycoplasmas*; in the bedbug *Cimex lectularius*, the Gram-negative Wolbachia can synthesize biotin and riboflavin which, are crucial for the host growth and reproduction (71, 72). Riboflavin biosynthesis is common for symbiotic associations and therefore occurs even in small and optimized genomes size like *Wolbachia* (~ 1.48 Mb) *and Mycoplasma* (0.51 ~1,38 Mb).

The small genomes sizes reduce the metabolic capabilities of *Mycoplasma*, although they can utilise sugars as a source of carbon and energy via glycolysis (Halbedel 2007). The simple sugar pentose phosphate pathway and genes encoding enzymes of the tricarboxylic acid cycle are absent from *Mycoplasma* genomes. So, sugars are transported by the phosphotransferase system which play a regulatory role via the regulatory function of HPr kinase/phosphorylase, which, was also annotated in the assembled MAG *Mycoplasma* in this study (**Supp. Table 1**). This carbon source is available in the epithelial tissues of hosts and its metabolism leads to the formation of hydrogen peroxide, the major virulence factor of several species of *Mollicutes* (Halbedel 2007). Up until this point, the functions detected in the *Mycoplasma* MAG, such as riboflavin, and phosphotransferase, support the argument for the strong symbiotic association of mycoplasmas with the host epithelial tissues, however, we were unable to find virulence factors in this study.

Also, it is reported that many *Mycoplasma* species can modify their surface antigenic molecules with high frequency (64, 73) which may likely play a key role in outmaneuvering the host immune system. This ability may generate phenotypic heterogeneity in colonising *Mycoplasma* populations and provide fitness benefits such as evasion of host immune responses and to adaptation to the environmental changes (73, 74). The majority of the *variable surface antigenic molecules* of mycoplasmas are lipoproteins (74–76), which, depending on the species, are encoded by single or multiple genes (64, 77). The expression of these lipoproteins, due to extensive antigenic variation, is thought to be a major factor for immune evasion, for example the P35 lipoprotein and its paralogs, which are distributed across the surface of *M. penetrans* cells, are immunodominant (78–80). Two lipoprotein encoding genes were found only in *Mycoplasma penetrans* but not in the *Mycoplasma*’s MAG in this study (Supp. Table 2).

Our study clearly demonstrates a potentially important ecological and functional association between *Mycoplasma* and *Salmo salar* that merits further investigation. This association probably reflects a mutualism rather than commensalism. Targeted meta-transcriptomics and strain-specific screening for this organism could improve our understanding of its role, biology, and function. Furthermore, targeted studies involving genome reduction and their association with the host dynamics are also necessary to fully understand the evolution of *Mycoplasma* symbiosis in *Salmo salar*. Further Omics investigations are also needed to assess the population genomics of mycoplasmas associated with Atlantic salmon to identify genetic variants in antibiotic resistant genes and the evolution of riboflavin pathway. New analytical genomic tools that can reveal further insights as well as bespoke experiments driven by the recommendations given in this study may possibly lead to the development of practices that can improve aquaculture industry especially whether we can demonstrate in the future a probiotic potential of mycoplasmas in salmonids.

## Acknowledgements

This research was supported in part by a research grant from Biotechnology and Biological Sciences Research Council (BBSRC) grant number BB/P001203/1, Science Foundation Ireland, the Marine Institute and the Dept. for the Economy, N. Ireland, under the Investigators Programme Grant No. SFI/15/IA/3028, and the Scottish Aquaculture Innovation Centre. The authors gratefully acknowledge the Glasgow Imaging and PolyOmics Facility for their support & assistance in this work’. Umer Zeeshan Ijaz is supported by NERC Independent Research Fellowship NERC NE/L011956/1 as well as Lord Kelvin Adam Smith Leadership Fellowship (Glasgow).

## Author Contributions

ML conceived and designed the study. PY, RK, MDN, TD, CH, PS, and EL collected the samples, performed FISH experiments, and analysed image data. BC performed the bioinformatics and interpreted the results. BC and ML wrote the original draft of the manuscript. All authors contributed to revisions of the manuscript.

## Legends of Supplementary Figures and Tables

**Supp. Figure S1.** Evaluation of the specificity of *Mycoplasma* probe against pure culture *Escherichia coli* and *Mycoplasma muris*. *Mycoplasma* probe Myc1-1 showed specific hybridization, giving a positive signal solely with *Mycoplasma muris*.

**Supp. Figure S2.1. The distribution of bacteria in farmed Salmon egg.** Very low abundance of bacteria was observed. Sections of egg samples were hybridized with Myc1-1 probe (Cy3, orange), Gam-1, FIR-1, EUB338, EUB338 II, EUB338 III probes (Cy5, red) and stained with DAPI (blue). **a)** Overlay of Cy3, Cy5, and DAPI filter set showed bright signal of *Mycoplasma* in Salmon egg, scaled at 10 μm. **b**) No signal of *Mycoplasma* was detected in egg samples, scaled at 50 μm.

**Supp. Figure S2.2. Distribution of *Mycoplasma* in the stomach of Salmon Parr.** (a) Image of DAPI signals (blue), (b) Gam-1, FIR-1, EUB338, EUB338 II, EUB338 III probes with Cy5 (red), (c) Image of *Mycoplasma* Specific probe Cy-3(orange), (d) Overlay image of all channels, (e) and (f), *Mycoplasma* signals were clustered in small groups. These are scaled at 100 μm for (a,b,c,d,) 20 μm for (e), and 10 μm for (f).

**Supp. Figure S2.3. Distribution of *Mycoplasma* in the Pyloric caecum of Salmon Parr**. The images were an overlay of DAPI signals (blue), hybridization signals of Gam-1, FIR-1, EUB338, EUB338 II, EUB338 III probes (Cy5, red) and *Mycoplasma* Specific Myc1-1 probe (Cy3, orange). (a) Signals of *Mycoplasma* aggregates on the muscularis mucosae, (b) epithelium of pyloric caecum. These are scaled at 10 μm for (a) and 10 μm for (b).

**Supp. Figure S2.4. Distribution of *Mycoplasma* in the midgut of Salmon Parr**. Overlay of all channels. **(a)**, *Mycoplasma* showed low abundance with small amount bacteria detected in midgut sections. (**b)**, Signals of *Mycoplasma* aggregated near epithelium cell nuclei. These are scaled at 20 μm for (a) and 10 μm for (b).

**Supp. Figure S2.5. Distribution of *Mycoplasma* in Stomach of adult Salmon**. (a), DAPI, (b), MycI probe, (c) all other bacteria probes, and (d) all layers overlaid. More signals were observed using universal probes in b than c using *Mycoplasma* specific probe. These are scaled at 50 μm.

**Supp. Figure S2.6. Distribution of *Mycoplasma* in Pyloric caecum of adult Salmon**. Signals of bacterial aggregates on the muscularis mucosae, lamina propria (**a**, arrows) and epithelium (**b**) of the pyloric caecum. **(c)** and (**d)**, overlay of all channels and differential interference contrast (DIC) image showed *Mycoplasma* signals were clustered in high abundance around epithelium cell nuclei. These are scaled at 50 μm for (a), 20 μm for (b), 5 μm for (c) and (d) respectively.

**Supp. Figure S2.7. Distribution of *Mycoplasma* in midgut of adult Salmon**. Overlay of all channels scaled at 20µm. (**a)**, *Mycoplasma* was distributed closely to epithelium cells. (**b)**, Overlay of all channels and DIC image confirmed *Mycoplasma* were in epithelium cells.

**Supp. Figure S3. Circular track highlighting the function** annotations of the *Mycoplasma* MAG from this study. **The functions are predicted on (a)** the negative strand 3’-5’ and (b) the positive strand 5’-3’.

**Supp. Figure S4.** Relationships of lifestyle and genome reduction. These include (a) plot of genes count and genome size, (b) plot of genes count and pseudogenes, and (c) the proportion of non-coding DNA across lifestyle.

**Supp. Table 1.** Summary of SEED subsystems annotations of the *Mycoplasm*a MAG from pyloric caecum of Atlantic salmon (*Salmo salar*)

**Supp. Table 2.** Summary of shared and unique SEED subsystems after pairwise genomic comparison of the *Mycoplasma* MAG from this study and *Mycoplasma Penetrans*

**Supp. Table 3.** Summary of best CDS orthologs based on best reciprocal BLAST hits between the *Mycoplasma* MAG in this study and *Mycoplasma Penetrans*

**Supp. Table 4.** Summary of genomic features from 247 strains of mycoplasmas. These data were collected from IMG database

## Supplementary files

**Supp. File 1.** The complete DNA sequence of the *Mycoplasma* MAG recovered from the pyloric caecum of the Atlantic salmon (*Salmo salar)*

**Supp. File 2.** Predicted coding regions (CDS) of genes in *Mycoplasma* MAG recovered from *Salmo salar*

**Supp. File 3.** Sequence alignment in (PHYLIP format) format of 137 16s genes from environmental and host-associated Mycoplasmas and Spiroplasmas.

**Supp. File 4**. Accession numbers and functions of the 21 markers concatenated in the MLST-based tree.

**Supp. File 5.** Sequence alignment (PHYLIP format) of 55 protein from the 21 concatenated from free-living and parasitic mycoplasmas

**Supp. File 6.** Sequence name abbreviation of tips labels within the 16s gene tree

**Supp. File 7.** Sequence name abbreviation of tips labels within the concatenated markers MLST-based tree

